# Snakeobjects: an object-oriented workflow management system

**DOI:** 10.1101/2022.12.08.519599

**Authors:** Boris Yamrom, Yoon-ha Lee, Steven Marks, Lubomir Chorbadjiev, Hannah V Meyer, Ivan Iossifov

**Affiliations:** Cold Spring Harbor Laboratory, Cold Spring Harbor NY; SeqPipe Ltd., Sofia, Bulgaria; New York Genome Center, New York, NY

## Abstract

Snakemake is one of the most popular workflow management systems, particularly in biological sciences. Snakemake workflows are highly portable, scalable, and transparent. Moreover, they enable the painless reproduction of published results and adaption to similar data processing and analysis projects. Here we present Snakeobjects, an extension of Snakemake that considerably simplifies the development of workflows and improves their readability and adaptability while preserving all the benefits of Snakemake that ensure the large and growing community of its users.

## Introduction

In declarative workflow platforms, like make (1), and Snakemake (2, 3), a workflow is a set of rules that define the transformation of a set of input files to a set of output/result files. Every rule lists the files it can produce, a procedure to determine the files it will use as input for a given result file, and the procedure to generate its result files from its input files. Notably, the input files for a rule can include files from the workflow input files or previously generated result files. For every resulting file, there must be one rule to generate it, but one rule can describe the generation of many files when the output files of the rule are listed as filename patterns. For example, when a Snakemake rule declares that it creates files with names like “`graph_{name}.png`,” it will be used to create all the results files whose names start with “`graph_`” and ends with `“.png`.” A workflow execution engine, like Snakemake, uses the workflow definition to generate the desired results files for a given set of input files. The engine applies the rules in order so that it executes a rule only after it has already generated all the input files the rule needs. An execution engine can run rules in parallel when it determines that the inputs and outputs of the rules are different.

Frequently workflows are large, with tens or even hundreds of rules that generate many files of many file types for a typical set of inputs. Large workflows, however, are associated with several difficulties. The workflow creators need to carefully organize the result files in a reasonable directory structure, making it easy for workflow users to find the files they need. They must also organize the rules across workflow definition files to allow for easy reading, understanding, maintaining, and reusing the workflow. While having all the rules in one workflow definition file in smallish workflows is acceptable, this becomes impractical for larger workflows. One of the most challenging tasks for the workflow developer is to create the procedures to identify the correct input files for a given result file. Specifying the necessary input files is easy for some rules (i.e., to generate a “`graph_{name}.png`” file, a rule may require the corresponding “`data_{name}.txt`” file), but it is not trivial for others. These procedures use the metadata and, in Snakemake, are usually implemented using small python lambda functions. As a result, the workflows become a mixture of metadata parsing sections, complex file naming schemas that are not completely clear when looking at one rule at a time, and instructions for creating the result files from the input files. That renders them challenging to develop, maintain, read and reuse.

Snakeobjects addresses these issues by grouping related result files into *objects*, organizing the objects into *object types* or sets of objects with similar result files, and interconnecting the objects into a dependency relationship encoded into an *object graph* that defines the order in which objects should be built. Taking advantage of this high-level organization leads to a clear workflow structure and substantially simplified rule definitions. Snakeobjects further enforces the separation of the *pipeline*, or the workflow definition, and the *project*, an application of the pipeline for a particular set of inputs. Snakeobjects creates a consistent directory structure within the project directory by creating a directory for each object and storing all the result files of an object (or the object’s *targets*) in that object directory. Throughout the presentation in the rest of the paper, we use the term pipeline instead of workflow in the context of Snakeobjects.

In the Results section below, we first describe Snakeobjects in the context of a non-trivial example shown in Figure 1. Then, in the Implementation section, we provide implementation details, and in the Discussion section, we address how Snakeobjects’ features improve the development, use, and maintenance of Snakemake workflows.

**Figure 1.**
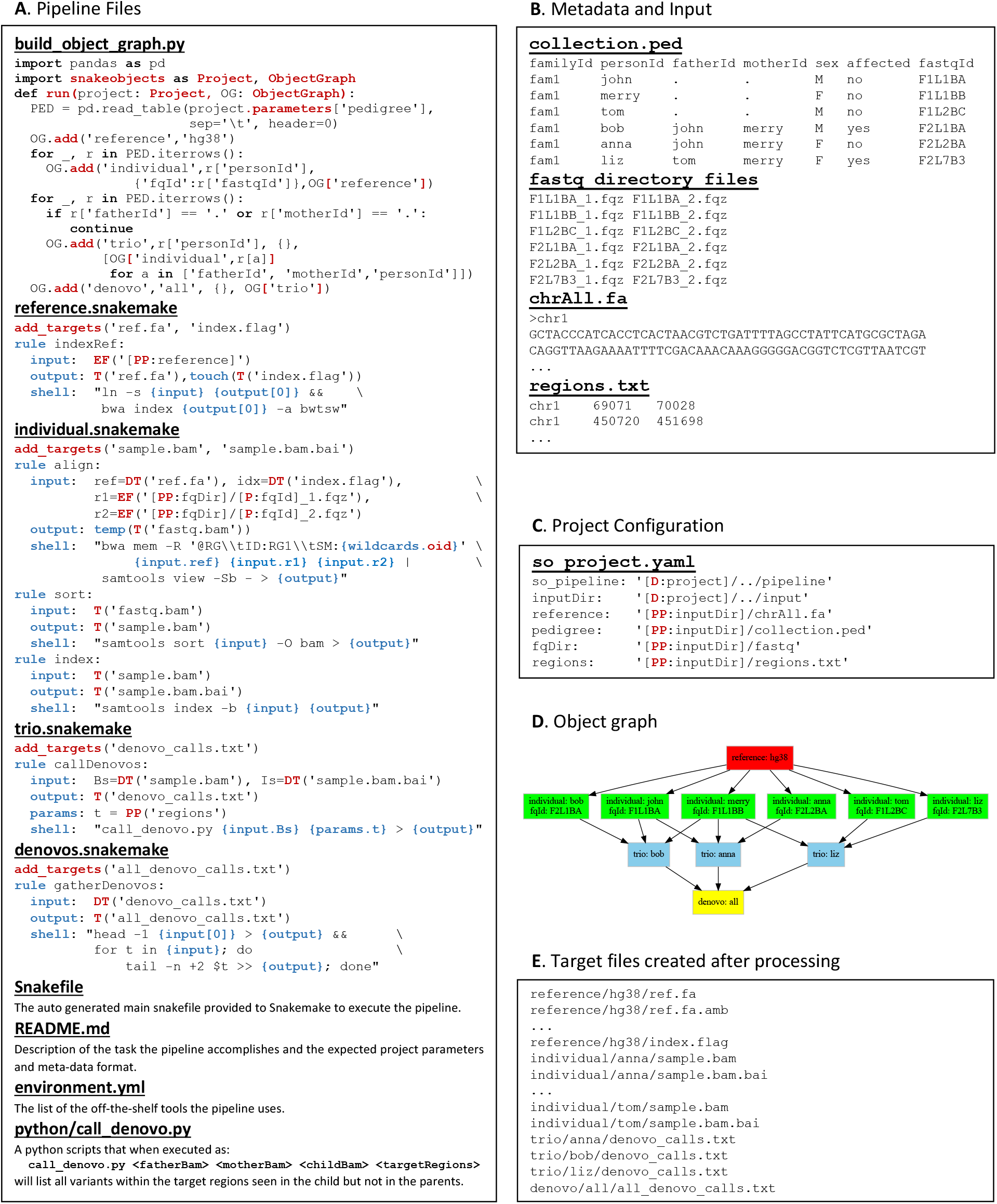
Example of a Snakeobjects pipeline and project. The figure demonstrates the use of Snakeobjects to design and use a pipeline for identifying de novo variants in a collection of families with FASTQ files provided for each family member. Panel **A** shows the pipeline’s files. The Snakeobjects and Snakemake specific components are colored red and blue, respectively. The **build_object_graph.py** script uses the metadata associated with a project to create a project object graph containing objects of one of four types: reference, individual, trio, and denovos. The pipeline includes a snakefile that lists the object-type targets and the rules to create them for each of the four object types. Python script call_denovo.py in callDenovos rule is an example of extra software tools used by the pipeline. The panels below show the metadata and input files for the project (**B**), the project’s configuration file (**C**), the object graph generated by the **build_object_graph.py** script for the project and drawn using the “sobjects graph” command (**D)**, and the resulting target files after successful execution of the pipeline (**E**).

## Results

### Description of the example

Panel A of Figure 1 shows a pipeline that identifies *de novo* substitutions in trios or families with a mother, a father, and a child. *De novo* substitutions are alleles/nucleotides at a given genomic position in the child that are not present in their parents at that position. As input, the pipeline uses new generation short sequencing reads stored in two FASTQ files for each involved individual. FASTQ files (4) are the standard file type produced by next-generation sequencing machines and often are produced in pairs when using pair-end sequencing mode. In addition to the files containing all identified *de novo* substitutions, the pipeline generates numerous intermediate files that may be of interest to the pipeline users, including reference genome index files, a BAM file (5), and a BAM index file for each individual containing the alignments of the individual’s FASTQ files to the reference genome, and a list of *de novo* substitutions identified in each trio.

Before we describe the example pipeline in detail, we will show what it accomplishes over three trios comprising six individuals (Figure 1, Panels B-E). Panel B shows the input and metadata files in this example application. The input comprises a directory called **fastq** containing the five pairs of FASTQ files for each individual, a reference genome sequence file (**chrAll.fa**), and a file containing the list of genomic regions (**regions.txt**) in which the *de novo* substitutions should be identified. The metadata has a form of a pedigree file (**collection.ped**), another standard file format used in genetics and genomics, that lists the individuals together with their names (the “`personId`” column), family relationships (the “`motherId`” and the “`fatherId`” columns), genders (“`sex`” column), affected statuses (“`affected`” column), and the prefix of the individuals’ FASTQ files (“`fastqId`” column). To apply a pipeline over particular inputs, the user creates a project, which is a directory with a project configuration file called **so_project.yaml**. The **so_project.yaml** file specifies the pipeline to be used (the “`so_pipeline`” property) and includes the project parameters that point to the input and the metadata (see Panel C). After the user prepares the project, they use Snakeobjects’ command line tool `sobjects` to execute the pipeline for the project. The pipeline stores the result files it generates in the project directory. Panel E shows a subset of the result files generated for our example.

### Object and object graph concepts

The core idea in Snakeobjects is to organize the result files into *objects* with closely related result files grouped into the same object and define a dependency relation between the objects through a directed acyclic *object graph*. For example, the pipeline in Figure 1 generates two files for every individual, a BAM file and a BAM index file. More realistic larger pipelines often generate many other files for each individual, like a VCF file (6) containing genetic variants, a gene or transcript level measure of RNA expression levels, or files containing quality measures. It is natural to group the files related to a given individual into an object.

In Snakeobjects, objects are characterized by an object type and an object id. In our example, we assign an object type `individual,` and object ids equal the individual’s `personId` (as specified in the project’s **collection.ped** file) to the objects corresponding to individuals. Objects of the same object type should have unique ids, but it is possible to have two objects of different object types have the same object id. Thus, to avoid confusion, we use the notation `<object type>/<object id>` to refer to objects. For example, the six individual objects are `individual/john`, `individual/merry`, `individual/tom`, `individual/bob`, `individual/anna`, and `individual/liz`.

We use the term *target* (borrowed from make (1)) to refer to the files generated for each object. The two targets generated for each `individual` object are `sample.bam` and `sample.bam.bai`. Every target within an object is associated with a result file, we call the target result file **<object type>/<object id>/<target>**. For example, the target result file for the target `sample.bam` of the object `individual/anna` is **individual/anna/sample.bam**. Snakeobjects stores the target results files in the project directory.

In addition to the `individual` objects, our pipeline uses one `reference` object to represent the result files related to the reference genome used in the processing, `trio` objects for each trio, and one `denovos` object that represents the overall list of the *de novo* substitutions. The object id for the `trio` objects is assigned to be the name of the `personId` of child of the trio, the id for the `reference` genome is set to `hg38`, and the id for the single `denovos` object is set to `all`. Objects may also be assigned object key-value parameters: the individual objects in our example are assigned the `fqId` parameter, whose values are the prefix of the FASTQ files related to the individual.

Finally, the objects are connected in an *object graph*, a directed acyclic graph whose edges specify dependencies between objects. An edge pointing from one object to another means that generating the targets of the second object requires the targets of the first object to be already generated. Panel D of Figure 1 shows the object graph for our example, with the object type, id, and parameters shown for each object represented with a rectangle and the dependency relationships represented with arrows between the rectangles. The rectangle’s color is determined based on the object type: red for `reference`, green for `individual`, light blue for `trio`, and yellow for the `denovos`.

### Snakeobjects pipelines

#### Core components

The Snakeobjects pipeline defines the procedure for creating the objects and the object graph for a project and the rules for generating the targets. The Snakeobjects rules are defined using targets in the context of an object graph and are simpler to write and understand than the equivalent Snakemake rules. The components of a Snakeobjects pipeline are stored in a pipeline directory. Every pipeline contains a **build_object_graph.py** file with the important job of creating the object graph for the project the pipeline operates on. **build_object_graph.py** is a python file with a function called `run` that accepts two parameters: `project` and `OG`, the object graph that will contain the project objects. The `run` function typically uses the `project` parameter to access the project’s meta-data, interpret it and add objects to the `OG` using `OG`’s `add` method, which has four arguments, specifying the new object’s type, id, parameters, and list of dependency objects. Importantly, the `OG` preserves the order of the dependencies, which is sometimes very helpful to pipeline developers. `OG` provides convenient methods for searching for objects that have already been added that can be used to assemble the list of dependent objects: “`OG[<object type>]`” returns the list of objects of type `<object type>` from the graph, and “`OG[<object type>,<object id>]`” returns the object of `<object type>` type and `<object id>` id. The top of Panel A of Figure 1 shows the **build_object_graph.py** script in our example pipeline that, although short, demonstrates all the important features.

Every pipeline must also contain one file named **<object type>.snakefile** for each object type used in the **build_object_graph.py** script. The pipeline directory in our example, thus, includes the four files, **reference.snakefile, individual.snakefile, trio.snakefile**, and **denovos.snakefile**, as shown on Panel A. Each of these **<object type>.snakefiles** contains the list of targets that will be generated for all objects of the object type (see the lines that use the `add_tareget` function calls) and the rules for creating these targets. Snakeobjects’ rules are written using Snakemake’s rule syntax and have four major components: the rule name, input definition, output definition, and procedure for generating outputs using the inputs. However, Snakeobjects differs from Snakemake in the way the outputs and inputs of the rules are specified.

#### Output specification and the T extension function

Outputs in Snakeobjects are specified using targets with the help of the extension function `T`, one of several extension functions provided by Snakeobjects. For example, the output definition “`output: T(‘sample.bam’)`” of the rule `sort` in the **individual.snakefile** file means that the rule will generate the `sample.bam` targets for every individual. When Snakeobjects uses a rule to create a target for an object, it will store the results in a file (target file) with a file name that has the form **<object type>/<object id>/<target>**. For example, when the target `sample.bam` is generated for the individual object with id `anna` (referred to as the `individual/anna` object), Snakeobjects will create the target file **individual/anna/sample.bam**. In contrast, the equivalent output definition in Snakemake, “`output: ‘individual/{oid}/sample.bam’,`” requires explicit specification of the complete file name, and every other rule that uses these files as input must specify the same full filename.

#### Input specification with EF, T, and DT extension functions

The inputs of Snakeobjects’ rules can be specified in three ways to allow for the three different sources of the input files. Input files for a rule can be from the project’s input files, targets of the current object (the object whose target the rule is generating), or targets of objects that the current object depends on. The `EF` extension function (`EF` stands for ‘external file’ and refers to files not created by the pipeline) can be used to access project input files. The `EF` function takes one string argument containing the input file’s path. Importantly, Snakeobjects applies its interpolation function over the provided string to allow the use of object and project parameters. The interpolation replaces sub-strings of the form `“[PP:<project parameter>]`” with the values of the `<project parameter>`s, as specified in the project configuration **so_project.yaml** file. Sub-strings of the form “`[P:<object parameter>]`” with the value of the `<object parameter>`s obtained from the current object’s representation in the object graph. `EF` function appears in two rules in our example pipeline. The input specification of `indexRef` rule in **reference.snakefile** is “`input: EF(‘[PP:reference]’)`” so that the rule will use the file pointed to by the `reference` parameter from the **so_object.yaml** file (see Panel C). The inputs of the `align` rule in **individual.snakefile** include “`EF(‘[PP:fqDir]/[P:fqId]_1.fqz’)`” and “`EF(‘[PP:fqDir]/[P:fqId]_2.fqz’)`”. These, when interpolated during the execution of the rule for the `individual/tom` object, result in **<project directory>/**..**/input/fastq/F1L2BC_1.fqz** and **<project directory>/**..**/input/fastq/F1L2BC_2.fqz** files, respectively, where **<project directory>** is the path to the project directory. Note that the interpolation also works for the values of the project parameters: the interpolation replaces the “`[D:project]`” substring with the project directory’s full path and the “`[PP:inputDir]`” with the value of the `inputDir` project parameter.

In addition to the output, the target extension function `T` can also appear in the input definitions of a rule, where it specifies that the rule will use the targets of the current objects as inputs. For example, when the `index` rule in **individual.snakefile** has both its input and output defined with the `T` function (“`input: T(‘sample.bam’)`” and “`output: T(‘sample.bam.bai’)`”). When Snakeobjects executes this rule for the `individual/bob` object, it will create the target file **individual/bob/sample.bam.bai** using the target file **individual/bob/sample.bam**.

Finally, the `DT` (or dependency target) extension function can specify inputs taken from targets of objects that the current object depends on, as specified in the object graph. In our example, all object types but the `reference` have one rule that uses the `DT` function. The `align` rule in **individual.snakefile** contains two `DT` calls, “`DT(‘ref.fa’)`” and “`DT(‘index.flag’)`”. When the rule is executed for any `individual` object, these are inferred to point to the **reference/hg38/ref.fa** and **reference/hg38/index.flag** target files, respectively, because all `individual` objects depend on the single `reference/hg38` object. The rule `callDenovos` in the **trio.snakefile** also has two calls of the `DT` function (“`DT(‘sample.bam’)`” and “`DT(‘sample.bam.bai’)`”). Still, the situation here is more complicated and interesting. When the rule is executed for the `trio/bob` object, “`DT(‘sample.bam’)`” returns the list of three target files: **individual/john/sample.bam, individual/merry/sample.bam**, and **individual/bob/sample.bam**. When the same rule is executed for the `trio/liz` object, “`DT(‘sample.bam’)`” returns a different list of target files: **individual/tom/sample.bam, individual/merry/sample.bam**, and **individual/liz/sample.bam**. These results become apparent when one examines the object graph shown in Panel D and the dependencies of the `trio` objects that point to the trio’s father, mother, and child (see the **build_object_graph.py** in Panel A). This rule takes advantage of the fact that the object graph maintains the order of the dependencies. Finally, the only `DT` call in the `gatherDenovos` rule in **denovos.snakefile** is “`DT(‘denovo_calls.txt’)`” and it returns the list of three target files, **trio/bob/denovo_calls.txt, trio/ann/denovo_calls.txt**, and **trio/liz/denovo_calls.txt**.

#### Additional pipeline components

In addition to the **build_object_graph.py** and the **<object type>.snakefile** files, a pipeline can also include several other components. These may consist of documentation files describing the pipeline’s task and providing instructions for the pipeline users; environment definition files that list the off-the-shelf packages the pipeline uses; and python, R, shell, or other types of scripts used in the pipeline’s rules. Our example includes three such components: **README.md, environment.yml**, and the **python/call_denovo.py** (see Figure 1, Panel A).

#### Using pipelines and projects

Users of a Snakeobjects pipeline typically follow three steps. First, they create a project directory and a **so_project.yaml** file in it that configures the project. Then, they build the object graph using the “`sobjects buildObjectGraph`” command. `sobjects` stores the resulting object graph in a file called **OG.json** in the project directory. Before continuing, the users can examine the object graph to ensure it is correct. They can use the “`sobjects describe`” command to obtain high-level statistics of the object graph (i.e., the number of objects for each object type) or the “`sobjects graph`” to generate a drawing of the object graph (like Panel D). The users can also directly examine the **OG.json**, which represents the graph in a simple, human-readable format. Finally, they use the “`sobjects run`” command to generate the target results files. The “`sobjects run`” command calls the Snakemake’s execution engine, thus preserving all the powerful Snakemake features.

### Implementation

#### Formal definition

Snakeobjects introduces an abstraction of workflows inspired by object-oriented design that replaces the low-level input-output relationships between files at the core of Snakemake’s rules. A *pipeline* (workflow) in Snakeobjects operates on *projects*, and projects are composed of *object*s. The objects within a project connect with dependency relationships organized in a directed acyclic graph called an *object graph*. Each object has a list of the objects it depends on (*dependency objects*). Each object also has a specified *object type*. Object types are characterized by *targets* that need to be created for each object of the given object type and the rules for creating the targets. The rules for building targets for an object type are included in a file named after the object type and are written using Snakemake’s syntax, where the inputs and outputs specify targets instead of files. Crucially, inputs can refer to targets of the current object and targets of the dependency objects of the current object. Finally, projects and objects can be associated with key-value parameters.

#### Pipelines and projects

Figure 2 shows the general structure of a project and a pipeline. The pipeline definition and the project data are usually kept in two separate directories. The project configuration file, **so_project.yaml**, is stored in the project directory and contains parameters that specify the pipeline operating on the project, pointers to the input and metadata associated with the project, and parameters that control the processing. The project directory also contains the **OG.json** file representing the project’s object graph and the target result files generated by the pipeline. The project metadata can have an arbitrary form (i.e., a list of input files, one or more CSV files, relation database, etc.) and is used to generate the project-specific object graph by a python script **build_object_graph.py** located in the pipeline directory. Snakeobjects provides the python class `ObjectGraph` that aids in the creation of the object graph.

**Figure 2.**
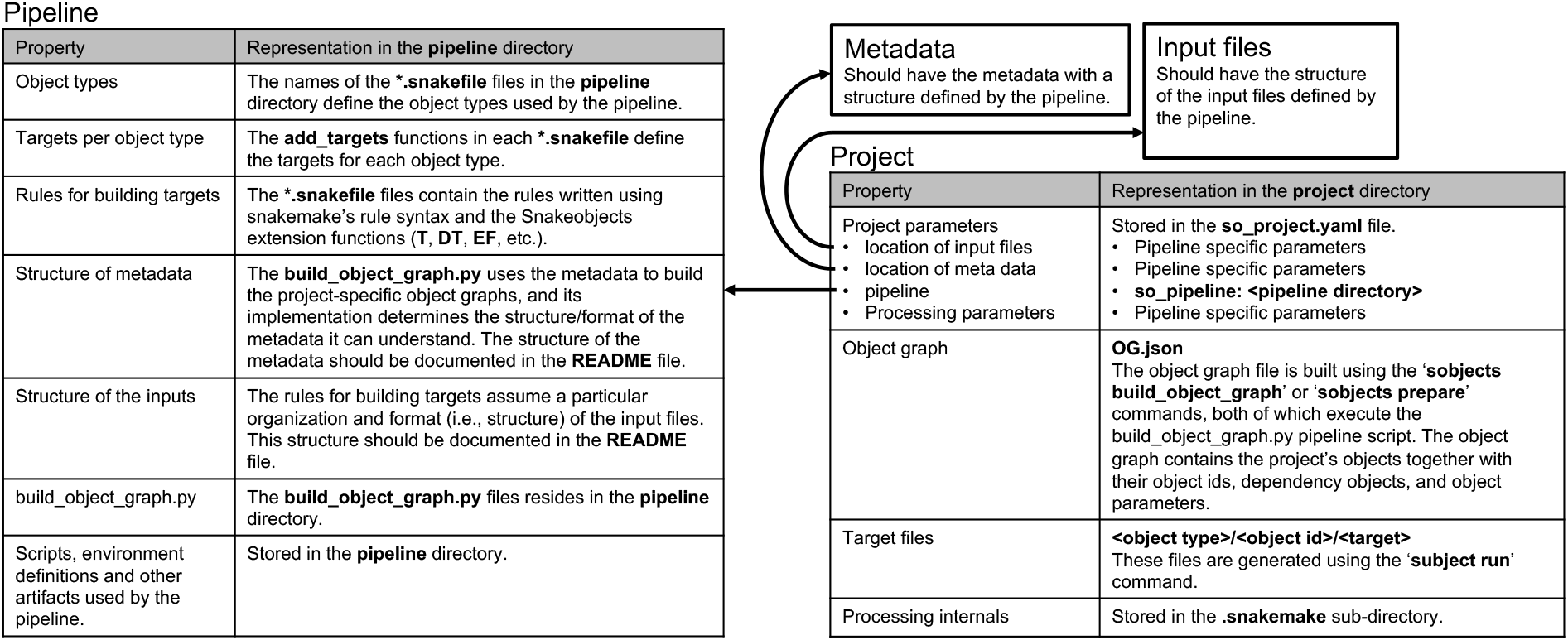
The general structure of pipeline and project directories.

The pipeline directory also contains one **<object type>.snakefile** file for each object type used in the pipeline. The **<object type>.snakefile** files use the Snakeobjects interpolation feature and a set of extension functions for referring to targets in the current object (`T`); to targets in dependency objects (`DT`); to parameters of the current object (`P`); to parameters of the dependency objects (`DP`); to project parameters (`PP`); and to external files (`EF`; most often the project’s input files). The pipeline directory can also contain scripts, environment definitions, or other artifacts used by the pipeline.

#### Main snakefile and Snakeobjects implicit targets

Snakeobjects operates through a file named **Snakefile** within the pipeline directory that the pipeline creator can generate with the “`sobjects createSnakefile”` command. We refer to this file as the main snakefile. The **Snakefile** imports the necessary Snakeobjects python classes and functions, instantiates a project object, and includes all the **<object type>.snakefile** files. The **Snakefile** also creates several implicit targets. For example, the **Snakefile** creates an `obj.flag` target for every object in the object graph. This target is dependent on all of the object’s targets and the `obj.flag` targets of all the object’s dependency objects. In addition, the **Snakefile** creates several global Snakemake targets: the `so_all_targets` is the default target and represents the complete set of project targets, and the `so_all_<object type>` represents all objects of type `<object type>.` The user can use these to generate subsets of the target result files. When the user executes “`sobjects run”` Snakeobjects will generate all the project’s target result files because the `so_all_targets` is the default target. When a user of our example from Figure 1 executes “`sobjects run trio/anna/denovos.txt,”` Snakeobjects will generate only the target results files necessary to generate the **trio/anna/denovos.txt** file. When the same user executes “`sobjects run individual/anna/obj.flag”,` Snakeobjects will generate all targets of the `individual/anna` object. Finally, if the user executes “`sobjects run so_all_individual”,` all targets of all `individual` objects will be generated.

#### snakeobjects python package

When installed, Snakeobjects provides a python package called `snakeobjects` that contains the `Project` and `ObjectGraph` classes. These classes are often used from the **<object type>.snakefile** or within the additional python scripts included in the pipeline. In addition, the `snakeobjects` includes the module `snakeUtils,` which contains all the extension functions described above and a few that we have not discussed here. It will be informative to advanced users to study the implementation of these functions.

#### Snakeobjects example and tutorial

We designed the example in Figure 1 to demonstrate Snakeobjects’ core concepts. The complete set of files and instructions needed to run the example is available as Supplementary Dataset 1, attached to this manuscript. Future updates will be posted at https://snakeobjects.readthedocs.io/en/stable/examples.html#snakeobjects-paper-example.As a comparison, we also included a pure Snakemake implementation of the example (see below for a discussion of the differences). Although it may be enough for experienced Snakemake users, the example and our presentation above are insufficient for new users to start using Snakeobjects. To guide new users, we created a lengthy Tutorial (https://snakeobjects.readthedocs.io/en/stable/tutorial.html) that shows the complete process of creating a complex pipeline, starting from the very beginning with all features explained in detail. In addition, the tutorial demonstrates how Snakeobjects handles evolving pipelines and growing projects (when the input data grows) gracefully.

## Discussion

Snakeobjects is an extension of Snakemake that improves several aspects of designing and using workflows by borrowing ideas from object-oriented design. Snakeobjects’ workflows are easier to write and read, are more modular, and are easier to adapt than the equivalent Snakemake pipelines. Specifically, Snakeobjects addresses the tedious tasks of organizing the project files and identifying the inputs for rules associated with all but the most straightforward Snakemake workflows. It is the Snakeobjects’ abstraction that enables these improvements: projects are comprised of objects of domain-specific object types, and objects are connected by dependency relationships encoded in an object graph. Importantly, workflows written in Snakeobjects are valid Snakemake workflows. Thus, Snakeobjects inherits all the Snakemake’s power.

Snakeobjects addresses two related problems that frequently occur in developing workflows with Snakemake. The first is that the workflow developer needs to design a directory structure to organize the workflow files. Even for medium-sized workflows, this becomes a non-trivial task. Snakeobjects implements a consistent file location structure using the concepts of object, object type, and targets of an object type and the extension function `T` that frees the developer from the chore of designing an arrangement themselves. The second problem is identifying the input files in workflows with complex object structures. The typical solution in Snakemake is lambda functions, python snippets that access a piece of metadata to identify inputs for a given rule. Unfortunately, such lambda functions are used in many of the rules, leading to copies of similar code dealing with specifics of the projects’ metadata repeated in many places in the workflow definitions or requiring developers to design project-specific approaches. Snakeobjects addresses the second problem using the object dependency graph, an object graph built outside the workflow definitions by the **build_object_graph.py** script that can be the only place with the knowledge of the project-specific metadata structure. The Snakemake rules are then cleaned of the clutter of numerous lambda functions and refer to inputs using the universal `T` and `DT` extensions functions to access the targets of the current object or its dependency objects following the dependency links in the objects graph.

To demonstrate how Snakeobjects simplifies the workflow definitions, we provide a pure Snakemake implementation in Supplementary Figure 1 of the pipeline presented in Figure 1. We created our Snakemake implementation by stripping the Snakeobjects’ features from the Snakeobjects implementation most straightforwardly. As a result, the Snakemake implementation creates identical result files using rules that are different from the Snakeobjects’ rules only in their input and output definitions. A quick comparison of the two implementations makes it clear that the Snakeobjects’ versions are substantially shorter, simpler, and thus clearer, with the difference being most drastic for the input sections of the `align`, `callDenovos`, and `gatherDenovos` rules. The main cause of the extra complexity is that the definitions in Snakemake directly manipulate the project’s metadata. Although such a Snakemake implementation style is typical (i.e., in small workflows or workflows created by beginner Snakemake users), Snakemake provides tremendous flexibility, and experienced Snakemake users would be able to improve our Snakemake implementation easily. But the improvements will be specifically crafted for the project, metadata, and input formats. In contrast, the Snakeobjects features, like object graph and extension functions, are general and available to simplify any workflow regardless of size.

But more than the solution or the specific technical issues, Snakeobjects pipelines benefit from the structure imposed by focusing on the high-level concepts of objects, object types, object graphs, pipelines, and projects. Objects are natural entities that the pipeline creator thinks about when designing a pipeline. Objects and their object types are usually clear from the start and provide a helpful scaffold for the pipeline where details, like the object targets, can be added further. The relationships between objects are also usually natural and can be defined early. Snakeobjects makes it easy to represent such relationships in an object graph, and the object graph allows for a much more natural definition of the complex rules for generating targets that depend on targets across dependency objects.

The benefits only grow when the size or the number of pipelines one deals with increases. With the consistent structure and the small set of universal extension functions mastered, it becomes easy to understand pipelines (like old pipelines created by the reader or someone else) and confidently extend them. Moreover, having all rules for an object type kept together in a separate **<object type>.snakemake** makes it easy to reuse object types across pipelines. After someone has access to a library of pipelines and object types (prepared by themselves, their colleagues, or others in a similar field), building new pipelines becomes a matter of copying the files that define the preexisting object types.

Although it is possible to have a Snakeobjects project and the pipeline that operates on it to be in the same directory, Snakeobjects sets a clear boundary between projects and pipelines. Such boundary ensures that the pipelines defined with Snakeobjects are general and can operate on different sets of inputs (or projects). Even in the typical situation when one creates a pipeline for one specific project, this separation comes handy: the user can create a small project on the subset of inputs that can be used for test and debug purposes (a strategy demonstrated in the Snakeobjects’ tutorial).

Snakeobjects is available as an open-source project (https://github.com/iossifovlab/snakeobjects) and is distributed as a Bioconda package (https://anaconda.org/bioconda/snakeobjects). The package is extensively documented (https://snakeobjects.readthedocs.io), and the documentation includes a tutorial that should enable both novices and Snakemake experts to create and use complex Snakeobjects workflows quickly.

## References

1. S. I. Feldman, Make — a program for maintaining computer programs. Software: Practice and Experience 9, 255–265 (1979).

2. J. Köster, S. Rahmann, Snakemake-a scalable bioinformatics workflow engine. Bioinformatics 34, 3600 (2018).

3. F. Mölder et al., Sustainable data analysis with Snakemake. F1000Res 10, 33 (2021).

4. P. J. Cock, C. J. Fields, N. Goto, M. L. Heuer, P. M. Rice, The Sanger FASTQ file format for sequences with quality scores, and the Solexa/Illumina FASTQ variants. Nucleic Acids Res 38, 1767–1771 (2010).

5. H. Li et al., The Sequence Alignment/Map format and SAMtools. Bioinformatics 25, 2078–2079 (2009).

6. P. Danecek et al., The variant call format and VCFtools. Bioinformatics 27, 2156–2158 (2011).

