## Supplementary Figure 1 for "Snakeobjects: an object-oriented workflow management system"

### Supplemental Figure 1. Pure Snakemake implementation of the example from Figure 1

```
import pandas as pd
import pathlib

input_dir = pathlib.Path("../input")
pedigree = input_dir / "collection.ped"
reference = input_dir / "chrAll.fa"
fq_dir = input_dir / "fastq"
regions = input_dir / "regions.txt"

PED = pd.read_table(pedigree, sep='\t', header=0)

rule all:
  input: 'denovos/all/all_denovo_calls.txt'

rule indexRef:
    input: reference.resolve()
    output: 'reference/all/ref.fa', touch('reference/all/index.flag')
    shell: "ln -s {input} {output[0]} &&     \
           bwa index {output[0]} -a bwtsw"

rule align:
  input:
    ref='reference/all/ref.fa', \
    idx='reference/all/index.flag', \
    r1=lambda wc: fq_dir / (PED[PED.personId == wc.oid]['fastqId'] + "_1.fqz"), \
    r2=lambda wc: fq_dir / (PED[PED.personId == wc.oid]['fastqId'] + "_2.fqz")
  output: temp('individual/{oid}/fastq.bam')
  shell: "bwa mem -R '@RG\\tID:RG1\\tSM:{wildcards.oid}' \
               {input.ref} {input.r1} {input.r2} |        \
         samtools view -Sb - > {output}"

rule sort:
  input: 'individual/{oid}/fastq.bam'
  output: 'individual/{oid}/sample.bam'
  shell: "samtools sort {input} -O bam > {output}"

rule index:
  input: 'individual/{oid}/sample.bam'
  output: 'individual/{oid}/sample.bam.bai'
  shell: "samtools index -b {input} {output}"


rule callDenovos:
  input:
    Bs=lambda wc: [f'individual/{PED[PED.personId == wc.oid][a].values[0]}/sample.bam'    \
                           for a in ["fatherId", "motherId", "personId"]],    \
    Is=lambda wc: [f'individual/{PED[PED.personId == wc.oid][a].values[0]}/sample.bam.bai'\
                           for a in ["fatherId", "motherId", "personId"]]
  output: 'trio/{oid}/denovo_calls.txt'
  params: t = regions
  shell:  "./call_denovo.py {input.Bs} {params.t} > {output}"


rule gatherDenovos:
  input:  expand('trio/{trio}/denovo_calls.txt', \
                trio=list(PED.query('motherId != "." and fatherId != "."')['personId']))
  output: 'denovos/all/all_denovo_calls.txt'
  shell: "head -1 {input[0]} > {output} &&      \
          for t in {input}; do                  \
              tail -n +2 $t >> {output}; done"
```

***Supplemental Figure1. Snakefile  Pure Snakemake implementation of the example from Figure 1.*** *The red highlighted input and output is different from the input and output in the Snakeobjects implementation.*
