## Supplementary Dataset 1 for "Snakeobjects: an object-oriented workflow management system": README.txt

Contents
++++++++

chrAll.fa - reference genome

genes.txt - gene models

targetRegions.txt - regions targeted by the exome capture

collection.ped - description of the families

All the gzipped fastq files are named using the following pattern:
fastq/<fastqid>_<read number>.fqz
