## Supplementary Dataset 1 for "Snakeobjects: an object-oriented workflow management system": build_object_graph.py

```
import pandas as pd
from snakeobjects import Project, ObjectGraph


def run(project: Project, OG: ObjectGraph):
    PED = pd.read_table(project.parameters['pedigree'],
                        sep='\t', header=0)
    OG.add('reference', 'hg38')
    for _, r in PED.iterrows():
        OG.add('individual', r['personId'],
                   {'fqId': r['fastqId']}, OG['reference'])
    for _, r in PED.iterrows():
        if r['fatherId'] == '.' or r['motherId'] == '.':
            continue
        OG.add('trio', r['personId'], {},
               [OG['individual', r[a]]
                for a in ['fatherId', 'motherId', 'personId']])
    OG.add('denovos', 'all', {}, OG['trio'])
```
