## Supplementary Dataset 1 for "Snakeobjects: an object-oriented workflow management system": denovos.snakefile

```
add_targets('all_denovo_calls.txt')
rule gatherDenovos:
  input:  DT('denovo_calls.txt')
  output: T('all_denovo_calls.txt')
  shell: "head -1 {input[0]} > {output} &&      \
          for t in {input}; do                  \
              tail -n +2 $t >> {output}; done"
```
