## Supplementary Dataset 1 for "Snakeobjects: an object-oriented workflow management system": individual.snakefile

```
add_targets('sample.bam', 'sample.bam.bai')
rule align:
  input:  ref=DT('ref.fa'), idx=DT('index.flag'),         \
          r1=EF("[PP:fqDir]/[P:fqId]_1.fqz"),              \
          r2=EF("[PP:fqDir]/[P:fqId]_2.fqz")
  output: temp(T('fastq.bam'))
  shell:  "bwa mem -R '@RG\\tID:RG1\\tSM:{wildcards.oid}' \
               {input.ref} {input.r1} {input.r2} |        \
           samtools view -Sb - > {output}"
rule sort:
  input:  T('fastq.bam')
  output: T('sample.bam')
  shell:  "samtools sort {input} -O bam > {output}"
rule index:
  input:  T('sample.bam')
  output: T('sample.bam.bai')
  shell:  "samtools index -b {input} {output}"
```
