## Supplementary Dataset 1 for "Snakeobjects: an object-oriented workflow management system": README.md

# Snakeobjects Paper Example version 1.0
## Pipeline description

This pipeline identifies *de novo* substitutions in trios from pair-end 
sequencing data in FASTQ format. It uses a trivial algorithm for the
detection of _de_ novo* substitutions implemented in the script 
**python/call_denovo.py**.

Requires pairs of FASTQ files stored 
in one directory named with the names having the form **\<prefix>_1.fq.tgz**
and **\<prefix>_2.fq.tgz** for the read 1 and read 2 FASTQ files, with the
related read 1 and read 2 files having the sample prefix. 

The pipeline uses a pedigree file describing the individuals, their family 
relationships, and file names for the FASTQ files related to the individual. 
The pedigree file should have the columns: "peronsId", "motherId", "fatherId",
and "fastqId", where the values in the  "fastqId" column contain the prefix 
for the pair of FASTQ files for the individual. A value of "." in "motherId" 
and "fatherId" columns indicates that the parent is not included in the 
dataset.

The pipeline also needs a reference genome file represented in FASTA format 
and a file containing the regions where the pipeline will search for *de novo*
substitutions, as needed for whole-exome sequencing projects. 
The file should have three columns showing the chromosome, the start, and
the end position. 
## Description of the required project parameters

The pipeline uses the following project parameters:

* **fqDir**, pointing to the directory with the FASTQ files;
* **pedigree**, pointing to the pedigree file;
* **reference**, pointing to the reference genome FASTA file;
* **regions**, pointing to the file with the regions.

## Description of the results

The final list of *de novo* substitutions will be stored in the 
**denovo/all/all_denovos.txt** files.

The pipeline will also create alignment BAM files and their indexes for each 
individual named:
**individual/\<personId>/sample.bam** and 
**individual/\<personId>/sample.bam.bai.**
