## Supplementary Dataset 1 for "Snakeobjects: an object-oriented workflow management system": reference.snakefile

```
add_targets('ref.fa', 'index.flag')
rule indexRef:
  input:  EF('[PP:reference]')
  output: T('ref.fa'),touch(T('index.flag'))
  shell:  "ln -s {input} {output[0]} &&    \
           bwa index {output[0]} -a bwtsw"
```
