## Supplementary Dataset 1 for "Snakeobjects: an object-oriented workflow management system": Snakefile

```
#snakeobjects 3.1.1
#Snakeobjects types: ['denovos', 'individual', 'reference', 'trio']
shell.prefix("set -euo pipefail; ")
from snakeobjects import Project 
from snakeobjects.snakeUtils import set_project, T, TE, DT, LFS, P, DP, PP, B, EF, add_targets, get_targets
from snakeobjects.remoteProjects import download_project_files_from_remote

# CLOUD related reorganization
import os,sys

if ("SO_CONTAINER" in os.environ and 
  "SO_REMOTE" in os.environ and
  os.environ["SO_CONTAINER"] == 'yes'):
    if ":" in os.environ["SO_REMOTE"]:
      provider, bucket = os.environ["SO_REMOTE"].split(":")
      download_project_files_from_remote(provider,bucket)
      os.environ['SO_PIPELINE'] = "workflow"
      os.environ['SO_PROJECT'] = "./" + bucket 
    fixPP = os.environ['SO_PIPELINE'] + "/fix_permissions.sh"
    if os.path.exists(fixPP):
      os.system("bash %s" % fixPP)

project = Project()
so_pipeline=project.get_pipeline_directory()

set_project(project)

rule so_all_targets:
  input:
    expand("{of}", of=project.get_all_object_flags())

include: "denovos.snakefile"
rule so_all_denovos:
  input:
    expand("{of}", of=project.get_all_object_flags("denovos"))

rule so_denovos_obj:
  input: get_targets("denovos")
  output: touch("denovos/{oid}/obj.flag")

include: "individual.snakefile"
rule so_all_individual:
  input:
    expand("{of}", of=project.get_all_object_flags("individual"))

rule so_individual_obj:
  input: get_targets("individual")
  output: touch("individual/{oid}/obj.flag")

include: "reference.snakefile"
rule so_all_reference:
  input:
    expand("{of}", of=project.get_all_object_flags("reference"))

rule so_reference_obj:
  input: get_targets("reference")
  output: touch("reference/{oid}/obj.flag")

include: "trio.snakefile"
rule so_all_trio:
  input:
    expand("{of}", of=project.get_all_object_flags("trio"))

rule so_trio_obj:
  input: get_targets("trio")
  output: touch("trio/{oid}/obj.flag")
```
