## Supplementary Dataset 1 for "Snakeobjects: an object-oriented workflow management system": trio.snakefile

```
add_targets('denovo_calls.txt')
rule callDenovos:
  input:  Bs=DT('sample.bam'), Is=DT('sample.bam.bai')
  output: T('denovo_calls.txt')
  params: t = PP('regions')
  shell:  "call_denovo.py {input.Bs} {params.t} > {output}"
```
