## Supplementary Dataset 1 for "Snakeobjects: an object-oriented workflow management system": so_project.yaml

```
so_pipeline: '[D:project]/../pipeline'
inputDir:    '[D:project]/../input'
reference:   '[PP:inputDir]/chrAll.fa'
pedigree:    '[PP:inputDir]/collection.ped'
fqDir:       '[PP:inputDir]/fastq'
regions:     '[PP:inputDir]/regions.txt'
```
