## Supplementary Dataset 1 for "Snakeobjects: an object-oriented workflow management system": call_denovo.py

```
#!/usr/bin/env python

import pysam
import sys
import pandas as pd
import numpy as np
from collections import Counter, defaultdict

dadBamF, momBamF, chlBamF, targetFile = sys.argv[1:]

AFS = [pysam.AlignmentFile(bmf) for bmf in [dadBamF, momBamF, chlBamF]]
smIds = [{x['SM'] for x in AF.header['RG']}.pop() for AF in AFS]

print("\t".join([f'{p}Id' for p in ['dad', 'mom', 'chl']]
                + ['chrom', 'pos', 'newAllele']
                + [f'{p}.{a}' for p in ['dad', 'mom', 'chl'] for a in "ACGT"]))
TT = pd.read_table(targetFile, sep='\t', names=['chr', 'beg', 'end'])
for ri, rgn in TT.iterrows():
    cntBuff = defaultdict(list)
    for AF in AFS:
        plps = AF.pileup(rgn['chr'], rgn['beg'], rgn['end'])
        for plp in plps:
            cnt = Counter([n.upper() for n in plp.get_query_sequences()])
            cntA = np.array([cnt[n] for n in 'ACGT'])
            cntBuff[plp.reference_pos].append(cntA)
    for pos, cntAs in cntBuff.items():
        if len(cntAs) != 3:
            continue
        trioCnt = np.vstack(cntAs)
        dpths = trioCnt.sum(axis=1)
        for dnvAI in range(4):
            if trioCnt[2, dnvAI] >= 3 and \
               trioCnt[0, dnvAI] == 0 and \
               trioCnt[1, dnvAI] == 0 and \
               dpths[0] >= 10 and dpths[1] >= 10:
                print("\t".join(map(str,
                                    smIds + [rgn['chr'], pos, "ACGT"[dnvAI]]
                                    + list(trioCnt.flatten()))))

for AF in AFS:
    AF.close()
```
